# Dissemination of Carbapenem-resistance and Plasmids-encoding Carbapenemases in Gram-negative Bacteria Isolated from India

**DOI:** 10.1101/2020.05.18.102434

**Authors:** Prasanth Manohar, Sebastian Leptihn, Bruno S. Lopes, Nachimuthu Ramesh

**Affiliations:** Antibiotic Resistance and Phage Therapy Laboratory, Department of Biomedical Sciences, School of Bioscience and Technology, Vellore Institute of Technology (VIT), Vellore-632014, Tamil Nadu, India; Zhejiang University-University of Edinburgh (ZJU-UoE) Institute, Zhejiang University, Haining, Zhejiang 314400, China and Department of Infectious Diseases, Sir Run Run Shaw Hospital, Zhejiang University School of Medicine, Hangzhou, China; University of Edinburgh Medical School, Biomedical Sciences, College of Medicine & Veterinary Medicine, The University of Edinburgh, 1 George Square, Edinburgh, EH8 9JZ, United Kingdom; School of Medicine, Medical Sciences and Nutrition, Medical Microbiology, University of Aberdeen, Aberdeen, AB25 2ZD UK

**Keywords:** Carbapenem, Gram-negative bacteria, Plasmid incompatibility grouping, Conjugative plasmid, Carbapenem-resistance genes

## Abstract

Carbapenem resistance in Gram-negative bacteria is an ongoing public-health problem of global dimensions leaving very few treatment options for severely infected patients. This study focuses on the dissemination of plasmid-borne carbapenemase genes in Gram-negative bacteria in Tamil Nadu, India. A total of 151 non-repetitive isolates belonging to 11 genera were collected from a diagnostic center in Tamil Nadu. *E. coli* (n=57) isolates were classified as, Enteropathogenic (n=12), Enteroaggregative (n=9), Enterohemorrhagic (n=8), Enterotoxigenic (n=3), Enteroinvasive (n=1) and unclassified *E. coli* (n=24). Of the 45 *Klebsiella* species, 14 were K1 whereas 11 were K2 serotype and in 20 *Klebsiella* serotype could not be determined. Other isolates (n=49) consisted of *P. aeruginosa*, *S. typhi*, *E. cloacae*, *A. baumannii*, *S. marcescens*, *A. xylosoxidans*, *P. mirabilis* and *E. meningoseptica*. Of the 151 isolates, 71% (n=107) and 68% (n=103) were found to be resistant to meropenem and imipenem respectively. The most prevalent beta-lactamase gene was *bla*_NDM-1_ (21%, 12/57) followed by *bla*_OXA-181_ (16%, 9/57), *bla*_GES-9_ (n=8), *bla*_OXA-23_ (n=7), *bla*_IMP-1_ (n=3), *bla*_GES-1_ (n=11) and *bla*_OXA-51_ (n=9). The unusual presence of *bla*_OXA-23_ was seen in *E. coli* (n=4), and *bla*_OXA-23_ and *bla*_OXA-51_ (IncA/C) in *K. pneumoniae* (n=3). Plasmid incompatibility (inc/rep) typing results showed that the plasmids carrying resistance genes (n=11) belonged to IncX, IncA/C, IncFIA-FIB and IncFIIA groups. *E. coli* and *K. pneumoniae* were able to transfer plasmid-borne carbapenemase via conjugation. This study highlights the prevalence of carbapenem resistance and the acquisition of plasmid-borne carbapenemase genes in Gram-negative bacteria highlighting the role of plasmid transfer in disseminating resistance.

## 1.0 Introduction

Antibiotic resistance is an emerging global health problem due to the injudicious use of antibiotics [1]. It is considered a major clinical and public health problem because of increasing bacterial resistance to most of the available antibiotics including penicillin, cephalosporins, carbapenems, and colistin [1]. World Health Organization (WHO) recently listed carbapenem-resistant *Acinetobacter baumannii*, *Pseudomonas aeruginosa* and Extended Spectrum Beta-Lactamases (ESBL) -producing *Enterobacteriaceae* as pathogens that are of critical importance [2]. Gram-negative bacteria (GNB) especially *Enterobacteriaceae* have developed resistance towards a broad spectrum of antibiotics responsible for significant mortality around the globe [3]. Carbapenems are considered as one of the last resort antibiotics against infections caused by multi-drug resistant GNB [4]. The emergence of carbapenem resistance especially in *Enterobacteriaceae* is a threat to the patients, particularly with complex infections, immunocompromised conditions and multiple diseases [5]. Because pathogens that are resistant to carbapenems often shows high resistance to other commonly used antibiotics that are often used for treatment, not only the mortality rates are high with increased hospital stay, but also huge medical expenditure placing emotional, economic and financial burden on families especially in resource limited countries [6].

The assessment of the rise in global antibiotic resistance has become very difficult due to the increasing rate of multi-drug resistance shown by pathogens with no proper harmonized surveillance systems in resource limiting countries [7]. Moreover, the co-existence of more than one carbapenem resistance gene with other genes like plasmid mediated AmpC, or plasmid mediated quinolone resistance has resulted in an increased acquisition of resistance among *Enterobacteriaceae* for community as well as hospital acquired infections [8,9]. The carbapenem-hydrolyzing oxacillinases (CHDL) are the major resistance mechanisms to carbapenems in *A. baumannii*. The first report of OXA-23 beta-lactamase in *A. baumannii* was from United Kingdom, in 1985 [10]. Later, OXA-23 was found to confer carbapenem-resistance in *A. nosocomialis* [11] and recently, it was reported in members of *Enterobacteriaceae* family [12–14]. In 1996, the first report of OXA-51 type beta-lactamase was from Argentina and at present, there are more than 150 variants of OXA-51 were reported globally [15]. These intrinsic enzymes in *A. baumannii* are naturally chromosomal-borne but rare cases of plasmid-borne genes are also reported [16]. Earlier, we reported the distribution of carbapenem and colistin resistance, and the role of integrons serving as the horizontal gene transfer agents in disseminating resistance among Gram-negative bacteria [17,18]. In the present study, molecular characterization of Gram-negative bacteria was performed and the role of interspecies plasmid transfer as evolutionary mechanism of carbapenem resistance was determined.

## 2.0 Materials and methods

### 2.1 Isolate collection and classification

During January 2015 and December 2016, a total of 151 Gram-negative bacterial isolates were collected from Hi-Tech diagnostic center in Chennai, Tamil Nadu, India. Bacteria were isolated from urine, blood, pus, bronchial secretion, cerebrospinal fluid, pulmonary secretion and bile fluid. The collected isolates were received at the Antibiotic Resistance and Phage Therapy Laboratory, VIT, Vellore, for further analyses. Genomic DNA was extracted from all the isolates using boiling lysis method [18]. Bacterial identification was carried out using VITEK identification system (bioMerieux) and 16S rRNA gene nucleotide sequence analysis using universal primers 27F and 1492R [18]. The PCR products were sequenced and identified to the species level using the BLASTN tool.

### 2.2 Antibiotic susceptibility testing and Minimal Inhibitory Concentration

Antibiotic resistance profiling was performed using the disk-diffusion method according to CLSI guidelines. The antibiotics used for this study were gentamicin (10 μg), amoxyclav (30 μg), cefotaxime (30 μg), ertapenem (10 μg), amikacin (30 μg), meropenem (10 μg), colistin (10 μg) and cefepime (30 μg). Minimum Inhibitory Concentration (MIC) was determined by broth micro-dilution method for meropenem and imipenem as described previously [18] and the results were interpreted according to CLSI guidelines [19].

### 2.3 Molecular analysis of E. coli pathotypes and Klebsiella serotypes

The *E. coli* pathotypes namely enteropathogenic *E. coli* (EPEC); enterohemorrhagic *E. coli* (EHEC); enterotoxigenic *E. coli* (ETEC); enteroaggregative *E. coli* (EAEC) and enteroinvasive *E. coli* (EIEC) were identified as described earlier [20]. The *Klebsiella* serotypes K1, K2 and K5 were determined using PCR primers and conditions as described earlier [21].

### 2.4 Molecular analysis of resistance-related genes

The isolates were screened for the presence of carbapenem resistance genes *bla*_NDM_, *bla*_OXA-48-like_, *bla*_KPC_, *bla*_IMP_ and *bla*_VIM_ [18]. A second multiplex PCR was also performed for *bla*_DIM_, *bla*_BIC_, *bla*_GIM_, *bla*_SIM_ and *bla*_AIM_ [22]. The *bla*_OXA-1_, *bla*_OXA-4_, *bla*_OXA-30_, *bla*_GES-1_, *bla*_GES-9_ and *bla*_GES-11_ were screened as described earlier [23]. The *bla*_OXA-23-like_, *bla*_OXA-24-_ like, *bla*_OXA-51-like_, *bla*_OXA-58-like_ were screened as described by Karunasagar et al. [24]. The PCR amplicons of the resistance genes were sequenced and genes were confirmed using NCBI BLASTN programme.

### 2.5 Plasmid isolation and plasmid incompatibility grouping

Plasmid isolation was performed for all the isolates harbouring resistance genes. The isolation of plasmid DNA was performed using HiPurA Plasmid DNA Miniprep Purification Kit (Himedia, India). Chromosomal DNA contamination was checked using the 16S rRNA primers as described earlier [25]. Plasmid incompatibility (*inc/rep*) typing (FIA, FIB, FIC, HI1, HI2, I1-Ig, L/M, N, P, W, T, A/C, K, B/O, X, Y, F, and FIIA replicons) was performed using multiplex PCR following the primers and PCR conditions as described by Carattoli et al. [26].

### 2.6 Conjugation studies

Representative carbapenem-resistant isolates harbouring plasmid-borne resistances (n=11) were subjected for conjugation using broth-mating method [18]. Briefly, the donor strain (strains carrying resistance genes) and the recipient strain (*E. coli* AB1157, Str^r^) were grown overnight, and mixed in 9:1 ratio each of donor and recipient. The cells were kept undisturbed for 6 hours at 37°C and plated on to antibiotic containing medium. The isolates which grew on both meropenem and streptomycin were considered as transconjugants. All the transconjugants were confirmed for the presence of respective carbapenem resistance genes using PCR.

## 3.0 Results

### 3.1 Bacterial classification

In this cross-sectional study, a total of 151 non-duplicate, Gram-negative bacteria belonging to 11 genera were studied which include *Escherichia coli* (n=57, 37.7%), *Klebsiella pneumoniae* (n=40, 26.4%), *Klebsiella oxytoca* (n=5, 3.3%), *Pseudomonas aeruginosa* (n=10, 6.6%), *Salmonella typhi* (n=8, 5.2%), *Enterobacter cloacae* (n=8, 5.2%), *Acinetobacter baumannii* (n=7, 4.6%), *Serratia marcescens* (n=5, 3.3%), *Achromobacter xylosoxidans* (n=5, 3.3%), *Proteus mirabilis* (n=5, 3.3%) and *Elizabethkingia meningoseptica* (n=1, 0.6%). Most of the isolates were isolated from urine 37% (56/151) and blood 28% (42/151) and from other sources such as pus (7%), bronchial secretion (2%), cerebrospinal fluid (1%), pulmonary secretion (1%), bile fluid (5%) and unknown (19%).

### 3.2 Antibiotic susceptibility studies

Table 1 summarizes the antibiotic susceptibility pattern of all the isolates tested against eight different antibiotics. MIC for meropenem showed that 107/151 (71%) isolates were resistant (fig.1), whereas 128 (84.7%) isolates were meropenem-resistant by the disk-diffusion method. For imipenem, 68% (n=103) were resistant by micro-broth dilution method whereas 83% (n=125) resistant by the disk-diffusion method. MIC_50_ and MIC_90_ values for meropenem were 16 mg/L and 8 mg/L respectively and for imipenem MIC_50_ = 8 mg/L and MIC_90_ = 8 mg/L.

**Table 1:**
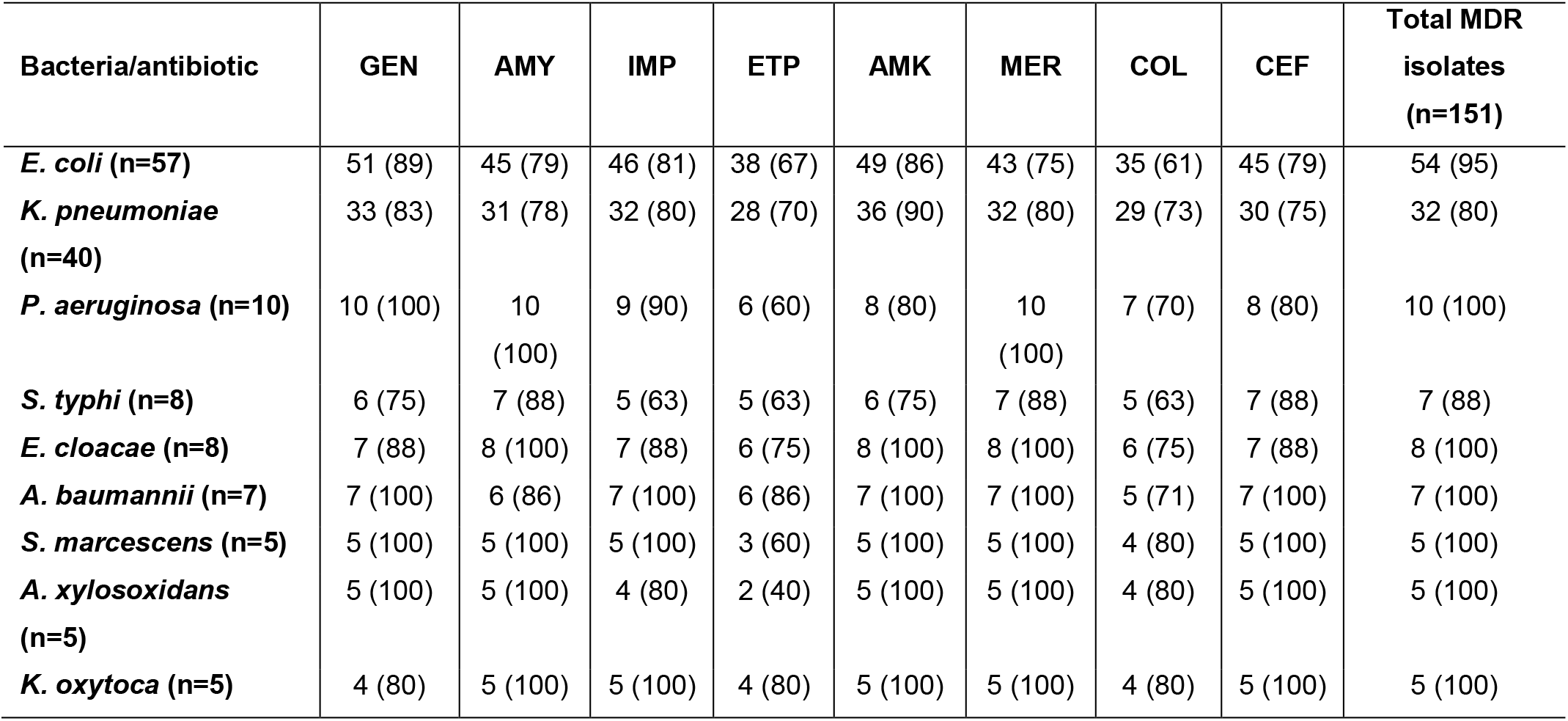
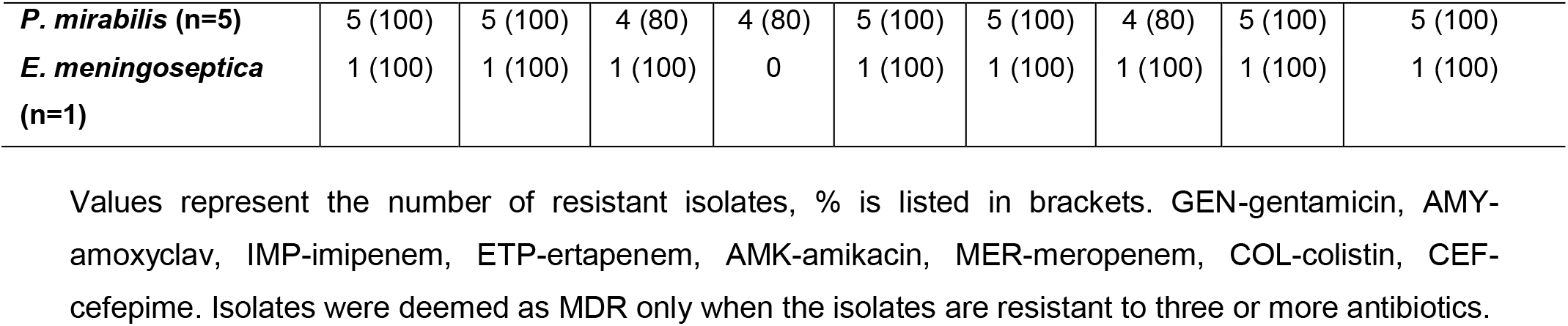
Antibiotic resistance pattern and the prevalence of multi-drug resistant isolates among 151 Gram-negative bacteria isolated from clinical samples.

| Bacteria/antibiotic | GEN | AMY | IMP | ETP | AMK | MER | COL | CEF | Total MDR isolates (n=151) |
| --- | --- | --- | --- | --- | --- | --- | --- | --- | --- |
| *E. coli* (n=57) | 51 (89) | 45 (79) | 46 (81) | 38 (67) | 49 (86) | 43 (75) | 35 (61) | 45 (79) | 54 (95) |
| *K. pneumoniae* (n=40) | 33 (83) | 31 (78) | 32 (80) | 28 (70) | 36 (90) | 32 (80) | 29 (73) | 30 (75) | 32 (80) |
| *P. aeruginosa* (n=10) | 10 (100) | 10 (100) | 9 (90) | 6 (60) | 8 (80) | 10 (100) | 7 (70) | 8 (80) | 10 (100) |
| *S. typhi* (n=8) | 6 (75) | 7 (88) | 5 (63) | 5 (63) | 6 (75) | 7 (88) | 5 (63) | 7 (88) | 7 (88) |
| *E. cloacae* (n=8) | 7 (88) | 8 (100) | 7 (88) | 6 (75) | 8 (100) | 8 (100) | 6 (75) | 7 (88) | 8 (100) |
| *A. baumannii* (n=7) | 7 (100) | 6 (86) | 7 (100) | 6 (86) | 7 (100) | 7 (100) | 5 (71) | 7 (100) | 7 (100) |
| *S. marcescens* (n=5) | 5 (100) | 5 (100) | 5 (100) | 3 (60) | 5 (100) | 5 (100) | 4 (80) | 5 (100) | 5 (100) |
| *A. xylosoxidans* (n=5) | 5 (100) | 5 (100) | 4 (80) | 2 (40) | 5 (100) | 5 (100) | 4 (80) | 5 (100) | 5 (100) |
| *K. oxytoca* (n=5) | 4 (80) | 5 (100) | 5 (100) | 4 (80) | 5 (100) | 5 (100) | 4 (80) | 5 (100) | 5 (100) |
| *P. mirabilis* (n=5) | 5 (100) | 5 (100) | 4 (80) | 4 (80) | 5 (100) | 5 (100) | 4 (80) | 5 (100) | 5 (100) |
| *E. meningoseptica* (n=1) | 1 (100) | 1 (100) | 1 (100) | 0 | 1 (100) | 1 (100) | 1 (100) | 1 (100) | 1 (100) |

Values represent the number of resistant isolates, % is listed in brackets. GEN-gentamicin, AMY-amoxyclav, IMP-imipenem, ETP-ertapenem, AMK-amikacin, MER-meropenem, COL-colistin, CEF-cefepime. Isolates were deemed as MDR only when the isolates are resistant to three or more antibiotics.

**Figure 1:**
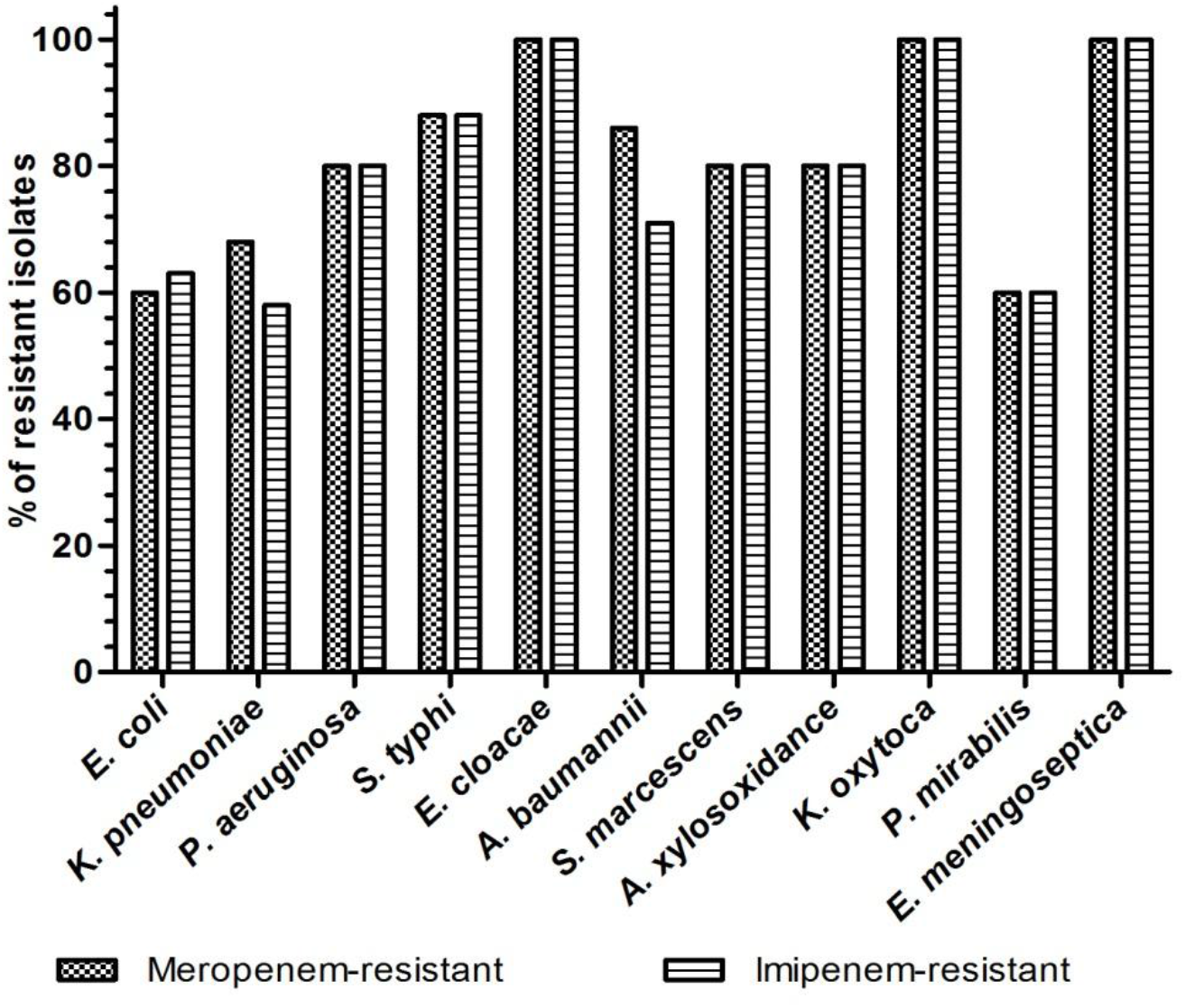
The distribution of Gram-negative bacteria and comparison of minimal inhibitory concentrations (MIC) of imipenem and meropenem resistance.

### 3.3 Distribution of carbapenemase resistance genes

The distribution of beta-lactamase resistance genes among 151 Gram-negative isolates is summarized in Table 2. Of the 57 *E. coli*, 32 isolates carried carbapenemase and five *E. coli* isolates carried more than one carbapenem resistance genes. Among the *K. pneumoniae*, 19/40 carried the studied genes and one isolate was positive for both *bla*_NDM_ and *bla*_OXA-48-like_. Carbapenem resistance genes were detected in 71/151 by PCR and 10 isolates had more than one gene. The most prevalent resistance gene was *bla*_NDM-1_ (n=22), *bla*_OXA-48-like_ (n=21), *bla*_GES-1_ (n=11), *bla*_GES-9_ (n=8), *bla*_OXA-23-like_ (n=7), *bla*_OXA-51-like_ (n=9) and *bla*_IMP-1_ (n=3). The beta-lactamase genes *bla*_KPC_, *bla*_VIM_, *bla*_BIC_, *bla*_GIM_, *bla*_DIM_, *bla*_SIM_ and *bla*_AIM_ were not detected in the isolates. Sequencing of genes showed that all the amplified NDM genes were NDM-1, OXA-48-like genes were OXA-181 and IMP genes were IMP-1.

**Table 2:** The distribution of carbapenemase genes among Gram-negative bacteria isolated from the clinical isolates. -A total of 20 resistance genes were studied that include *bla*_NDM_, *bla*_OXA-48-like_, *bla*_KPC_, *bla*_IMP_, *bla*_VIM_, *bla*_DIM_, *bla*_BIC_, *bla*_GIM_, *bla*_SIM_, *bla*_AIM_ *bla*_OXA-1_, *bla*_OXA-4_, *bla*_OXA-30_, *bla*_GES-1_, *bla*_GES-9_, *bla*_GES-11_, *bla*_OXA-23-like_, *bla*_OXA-24-like_, *bla*_OXA-51-like_, *bla*_OXA-58-like_. -The genes *bla*_KPC_, *bla*_VIM_, *bla*_DIM_, *bla*_BIC_, *bla*_GIM_, *bla*_SIM_, *bla*_AIM_, *bla*_OXA-1_, *bla*_OXA-4_, *bla*_OXA-30_, *bla*_GES-11_, *bla*_OXA-24-_ like and *bla*_OXA-58-like_ were not observed in any of the isolates. -GES-1 is an extended spectrum beta-lactamase which shows a significant degree of inhibition by imipenem indicating that it may be able to bind imipenem with storng affinity without being able to hydrolyze it. This enzyme was detected in 11 isolates. -OXA-51 is a beta-lactamase which can only act as a carbapenemase if it is upregulated by insertion elements or has a Trp222 site mutation. This enzyme was detected in 9 isolates.

| Bacteria/resistance gene | *bla*<sub>NDM-1</sub> | *bla*<sub>OXA-181</sub> | *bla*<sub>IMP-1</sub> | *bla*<sub>GES-1</sub> | *bla*<sub>GES-9</sub> | *bla*<sub>OXA-23</sub> | *bla*<sub>OXA-51</sub> |
| --- | --- | --- | --- | --- | --- | --- | --- |
| *E. coli* (n=57) | 12 | 9 | 1 | 6 | 5 | 4 | - |
| *K. pneumoniae* (n=40) | 5 | 7 | - | 3 | 2 | 1 | 2 |
| *P. aeruginosa* (n=10) | 1 | - | - | 2 | - | - | - |
| *E. cloacae* (n=8) | 2 | 1 | 1 | - | 1 | - | - |
| *A. baumannii* (n=7) | - | 2 | - | - | - | 2 | 7 |
| *S. marcescens* (n=5) | 1 | - | - | - | - | - | - |
| *A. xylosoxidans* (n=5) | 1 | - | - | - | - | - | - |
| *K. oxytoca* (n=5) | - | 1 | - | - | - | - | - |
| *P. mirabilis* (n=5) | - | 1 | 1 | - | - | - | - |
| **Total** | **22** | **21** | **3** | **11** | **8** | **7** | **9** |

### 3.4 Identification of E. coli pathotypes and Klebsiella serotypes

The *E. coli* pathotyping results showed that, of the 57 *E. coli* isolates tested 12 were Enteropathogenic (EPEC), 9 Enteroaggregative (EAEC), 8 Enterohemorrhagic (EHEC), 3 Enterotoxigenic (ETEC), 1 Enteroinvasive (EIEC) and 24 unclassified *E. coli* (Table 3). Of the 12 EPEC isolates 3 were positive for NDM-1, 2 for OXA-181 and one for OXA-23; among 8 EHEC isolates NDM-1 was detected in 2 isolates, OXA-181 in one isolate and GES-1 plus GES-9 in one isolate; among 9 EAEC isolates GES-1 was found in 1 isolate, OXA-23 in 1 and OXA-23 along with GES-9 in one isolate; among 3 ETEC isolates 1 isolate carried NDM-1 and one EIEC isolate carried NDM-1 among with OXA-181. The virulence genes found in *E. coli* isolates included *eaeA* (n=20), LT (n=3), *aggR* (n=6), *astA* (n=5) and *VirA* (n=1). 24 *E. coli* isolates did not belong to any of the tested pathotypes.

**Table 3:**
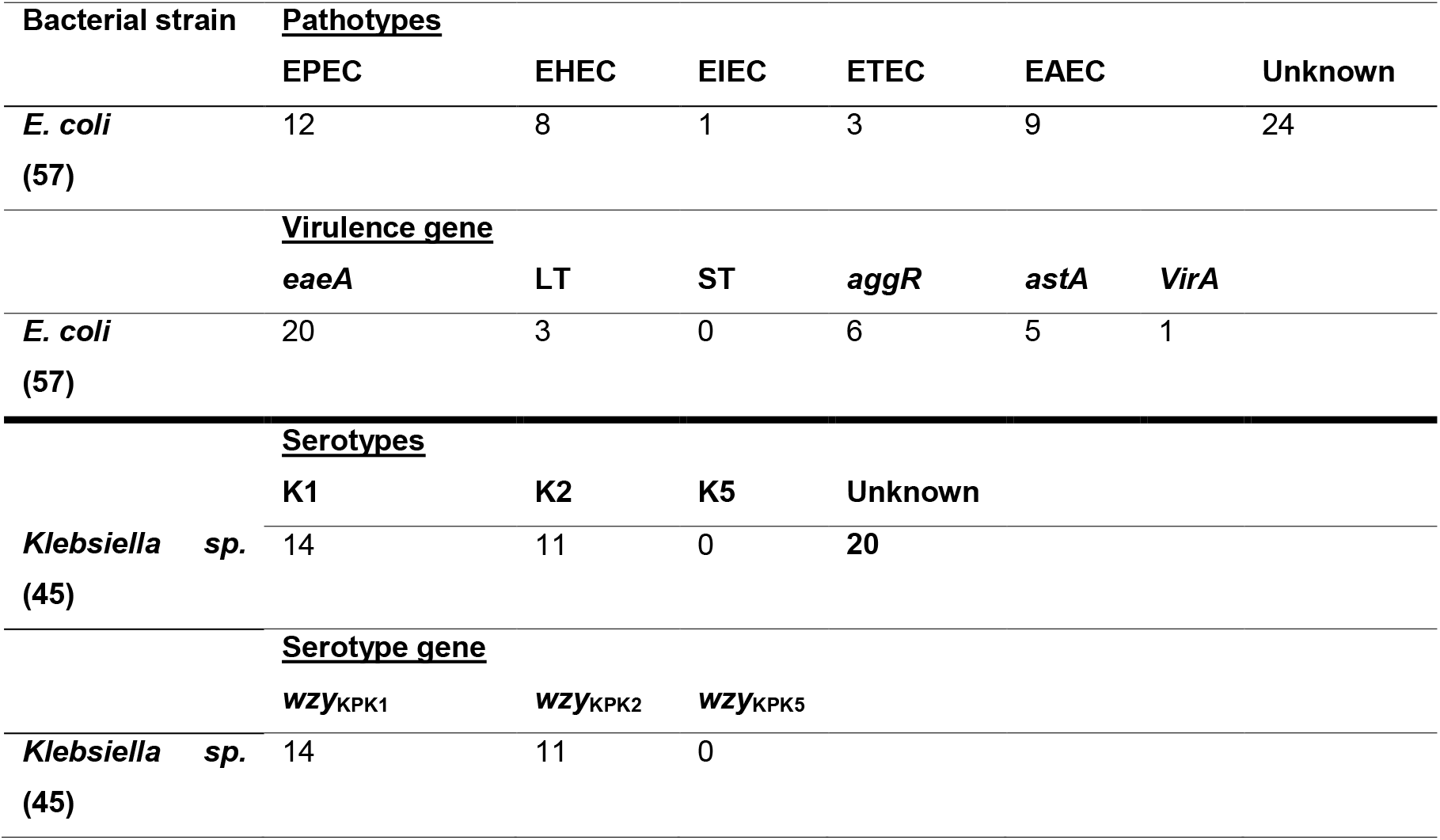
Number of *E. coli* pathotypes and *Klebsiella* serotypes among all the *E. coli* and *Klebsiella* species isolated from clinical samples.

| Bacterial strain | Pathotypes |  |  |  |  |  |
| --- | --- | --- | --- | --- | --- | --- |
|  | EPEC | EHEC | EIEC | ETEC | EAEC | Unknown |
| *E. coli*<br>(57) | 12 | 8 | 1 | 3 | 9 | 24 |
|  | Virulence gene |  |  |  |  |  |
|  | eaeA | LT | ST | aggR | astA | VirA |
| *E. coli*<br>(57) | 20 | 3 | 0 | 6 | 5 | 1 |
|  | Serotypes |  |  |  |  |  |
|  | K1 | K2 | K5 | Unknown |  |  |
| *Klebsiella* sp.<br>(45) | 14 | 11 | 0 | 20 |  |  |
|  | Serotype gene |  |  |  |  |  |
|  | wzy<sub>KPK1</sub> | wzy<sub>KPK2</sub> | wzy<sub>KPK5</sub> |  |  |  |
| *Klebsiella* sp.<br>(45) | 14 | 11 | 0 |  |  |  |

Of the 45 *Klebsiella* species, 14 belonged to K1 serotype, 11 were K2, none of the isolates were of K5 serotypes and 20 were of unknown serotypes (Table 3). Of the 14 *K. pneumoniae* K1, NDM-1 (n=1), OXA-181 (n=4) and OXA-51 (n=1) was detected in K1 serotypes. Among 11 *K. pneumoniae* K2, OXA-181 (n=1) and GES-9 (n=1) were detected in K2 serotypes.

### 3.5 Plasmid incompatibility typing and conjugation

Plasmid DNA was isolated from 70 isolates which carried resistance genes (Table 4). The plasmids ranged from 10-100 kb in size. In total, of the 151 isolates studied 70 isolates carried resistance genes of which 11 were plasmid-borne and 59 were chromosomal. Of the 37 *E. coli* isolates, 32 isolates carried resistance genes of which 6 were plasmid-borne. Among 40 *K. pneumoniae,* only 19 carried resistance genes of which 3 were associated with plasmids. In *E. cloacae*, one isolate carried *bla*_NDM-1_ on plasmid and in *P. mirabilis*; one isolate carried plasmid-borne *bla*_IMP-1_. Plasmid incompatibility/replicon (inc/rep) typing results showed that the plasmids were belonging to Inc/rep types: IncX, IncA/C, IncFIA-FIB and IncFIIA (Table 4). *E. coli* isolates harboured *bla*_NDM-1_ genes in IncX (EC10), IncA/C (EC21) and IncFIA-FIB (EC29) type plasmids whereas *bla*_OXA-48-like_ genes were associated with IncFIIA (EC39) and IncFIA-FIB (EC29), and *bla*_GES-1/9_ genes with IncFIA-FIB (EC47) type plasmid.

**Table 4:** Distribution of resistance genes, plasmid incompatibility grouping and transconjugation studies on Gram-negative isolates that were harbouring resistance genes.

| Bacterial isolate | Source | Pathotype/<br>Serotype | Resistance<br>gene | Plasmid<br>*inc/rep* typing | Conjugative<br>plasmid |
| --- | --- | --- | --- | --- | --- |
| *E. coli* EC1 | Urine | EPEC | *bla*<sub>NDM-1</sub> | - | - |
| *E. coli* EC2 | Urine | EAEC | ND | - | - |
| *E. coli* EC3 | Blood | EHEC | ND | - | - |
| *E. coli* EC4 | Pus | Unknown | *bla*<sub>NDM-1</sub> | - | - |
| *E. coli* EC5 | Urine | Unknown | *bla*<sub>NDM-1</sub> | - | - |
| *E. coli* EC6 | Pus | EAEC | ND | - | - |
| *E. coli* EC7 | Urine | EPEC | *bla*<sub>NDM-1</sub> | - | - |
| *E. coli* EC8 | Urine | Unknown | ND | - | - |
| *E. coli* EC9 | Urine | EAEC | ND | - | - |
| *E. coli* EC10 | Blood | EHEC | *bla*<sub>NDM-1</sub> | *IncX | + |
| *E. coli* EC11 | Unknown | Unknown | ND | - | - |
| *E. coli* EC12 | Urine | Unknown | *bla*<sub>NDM-1</sub> | - | - |
| *E. coli* EC13 | Unknown | Unknown | ND | - | - |
| *E. coli* EC14 | Unknown | EPEC | ND | - | - |
| *E. coli* EC15 | Urine | ETEC | ND | - | - |
| *E. coli* EC16 | Urine | Unknown | ND | - | - |
| *E. coli* EC17 | Urine | Unknown | *bla*<sub>NDM-1</sub> | - | - |
| *E. coli* EC18 | Urine | EPEC | ND | - | - |
| *E. coli* EC19 | Unknown | ETEC | ND | - | - |
| *E. coli* EC20 | Urine | Unknown | ND | - | - |
| *E. coli* EC21 | Blood | EHEC | *bla*<sub>NDM-1</sub> | *IncA/C | + |
| *E. coli* EC22 | Unknown | Unknown | *bla*<sub>NDM-1</sub> | - | - |
| *E. coli* EC23 | Urine | ETEC | *bla*<sub>NDM-1</sub> | - | - |
| *E. coli* EC24 | Unknown | EAEC | ND | - | - |
| *E. coli* EC25 | Urine | EPEC | *bla*<sub>NDM-1</sub> | - | - |
| *E. coli* EC26 | Urine | Unknown | ND | - | - |
| *E. coli* EC27 | Urine | EPEC | ND | - | - |
| *E. coli* EC28 | Bile fluid | Unknown | ND | - | - |
| *E. coli* EC29 | Unknown | EIEC | *bla*<sub>NDM-1</sub>,<br>*bla*<sub>OXA-181</sub> | *IncFIA-FIB | + |
| *E. coli* EC30 | Urine | Unknown | *bla*<sub>OXA-181</sub> | - | - |
| *E. coli* EC31 | Unknown | EPEC | *bla*<sub>OXA-181</sub> | - | - |
| *E. coli* EC32 | Blood | EAEC | ND | - | - |
| *E. coli* EC33 | Blood | Unknown | *bla*<sub>OXA-181</sub> | - | - |
| *E. coli* EC34 | Bile fluid | EPEC | *bla*<sub>OXA-181</sub> | - | - |
| *E. coli* EC35 | Urine | Unknown | ND | - | - |
| *E. coli* EC36 | Urine | Unknown | *bla*<sub>OXA-181</sub> | - | - |
| *E. coli* EC37 | Bile fluid | EPEC | ND | - | - |
| *E. coli* EC38 | Urine | EPEC | ND | - | - |
| *E. coli* EC39 | Blood | Unknown | *bla*<sub>OXA-181</sub> | *IncFIIA | + |
| *E. coli* EC40 | Blood | EHEC | *bla*<sub>OXA-181</sub> | - | - |
| *E. coli* EC41 | Blood | Unknown | *bla*<sub>OXA-181</sub> | - | - |
| *E. coli* EC42 | Blood | Unknown | *bla*<sub>IMP-1</sub> | - | - |
| *E. coli* EC43 | Urine | EPEC | ND | - | - |
| *E. coli* EC44 | Pus | EAEC | *bla*<sub>GES-1</sub> | *IncFIA-FIB | + |
| *E. coli* EC45 | Unknown | Unknown | *bla*<sub>GES-1</sub> | - | - |
| *E. coli* EC46 | Urine | EAEC | *bla*<sub>GES-1</sub> | - | - |
| *E. coli* EC47 | Blood | EHEC | *bla*<sub>GES-1</sub>, *bla*<sub>GES-9</sub> | *IncFIA-FIB | + |
| *E. coli* EC48 | Pus | Unknown | ND | - | - |
| *E. coli* EC49 | Blood | EHEC | *bla*<sub>GES-1</sub>, *bla*<sub>GES-9</sub> | - | - |
| *E. coli* EC50 | Pus | Unknown | *bla*<sub>GES-1</sub>, *bla*<sub>GES-9</sub> | - | - |
| *E. coli* EC51 | Bile fluid | Unknown | *bla*<sub>GES-9</sub> | - | - |
| *E. coli* EC52 | Unknown | EAEC | *bla*<sub>GES-9</sub>,<br>*bla*<sub>OXA-23</sub> | - | - |
| *E. coli* EC53 | Blood | Unknown | *bla*<sub>OXA-23</sub> | - | - |
| *E. coli* EC54 | Urine | EPEC | *bla*<sub>OXA-23</sub> | - | - |
| *E. coli* EC55 | Unknown | EHEC | ND | - | - |
| *E. coli* EC56 | Urine | EAEC | *bla*<sub>OXA-23</sub> | - | - |
| *E. coli* EC57 | Blood | EHEC | ND | - | - |
| *K. pneumoniae* KP1 | Urine | K1 | ND | - | - |
| *K. pneumoniae* KP2 | Urine | K2 | ND | - | - |
| *K. pneumoniae* KP3 | Blood | Unknown | *bla*<sub>NDM-1</sub> | - | - |
| *K. pneumoniae* KP4 | Bile fluid | K1 | ND | - | - |
| *K. pneumoniae* KP5 | Urine | K2 | ND | - | - |
| *K. pneumoniae* KP6 | Blood | K1 | ND | - | - |
| *K. pneumoniae* KP7 | Blood | Unknown | *bla*<sub>NDM-1</sub> | - | - |
| *K. pneumoniae* KP8 | Blood | K1 | ND | - | - |
| *K. pneumoniae* KP9 | Urine | K1 | *bla*<sub>NDM-1</sub> | - | - |
| *K. pneumoniae* KP10 | Blood | Unknown | *bla*<sub>NDM-1</sub> | *IncFIA-FIB | + |
| *K. pneumoniae* KP11 | Unknown | Unknown | *bla*<sub>NDM-1</sub> | - | - |
| *K. pneumoniae* KP12 | Bile fluid | K2 | ND | - | - |
| *K. pneumoniae* KP13 | Urine | K1 | ND | - | - |
| *K. pneumoniae* KP14 | Urine | Unknown | ND | - | - |
| *K. pneumoniae* KP15 | Pulmonary secretion | K2 | ND | - | - |
| *K. pneumoniae* KP16 | Urine | K2 | ND | - | - |
| *K. pneumoniae* KP17 | Blood | K1 | *bla*<sub>OXA-181</sub> | - | - |
| *K. pneumoniae* KP18 | Unknown | K2 | ND | - | - |
| *K. pneumoniae* KP19 | Blood | Unknown | *bla*<sub>OXA-181</sub> | - | - |
| *K. pneumoniae* KP20 | Unknown | K1 | *bla*<sub>OXA-181</sub> | - | - |
| *K. pneumoniae* KP21 | Unknown | Unknown | *bla*<sub>OXA-181</sub> | - | - |
| *K. pneumoniae* KP22 | Unknown | Unknown | ND | - | - |
| *K. pneumoniae* KP23 | Blood | K1 | ND | - | - |
| *K. pneumoniae* KP24 | Blood | K2 | ND | - | - |
| *K. pneumoniae* KP25 | Unknown | K2 | *bla*<sub>OXA-181</sub> | - | - |
| *K. pneumoniae* KP26 | Blood | Unknown | ND | - | - |
| *K. pneumoniae* KP27 | Unknown | K1 | *bla*<sub>OXA-181</sub> | - | - |
| *K. pneumoniae* KP28 | Blood | K1 | *bla*<sub>OXA-181</sub> | - | - |
| *K. pneumoniae* KP29 | Unknown | Unknown | ND | - | - |
| *K. pneumoniae* KP30 | Urine | K1 | ND | - | - |
| *K. pneumoniae* KP31 | Blood | Unknown | *bla*<sub>GES-1</sub> | *IncA/C | - |
| *K. pneumoniae* KP32 | Unknown | Unknown | *bla*<sub>GES-1</sub> | - | - |
| *K. pneumoniae* KP33 | Urine | Unknown | *bla*<sub>GES-1</sub> | - | - |
| *K. pneumoniae* KP34 | Blood | K2 | ND | - | - |
| *K. pneumoniae* KP35 | Unknown | K1 | ND | - | - |
| *K. pneumoniae* KP36 | Urine | Unknown | *bla*<sub>GES-9</sub> | - | - |
| *K. pneumoniae* KP37 | Bile fluid | K2 | *bla*<sub>GES-9</sub> | - | - |
| *K. pneumoniae* KP38 | Blood | K2 | ND | - | - |
| *K. pneumoniae* KP39 | Urine | Unknown | *bla*<sub>OXA-23,</sub><br>*bla*<sub>OXA-51</sub> | *IncA/C | - |
| *K. pneumoniae* KP40 | Urine | K1 | *bla*<sub>OXA-51</sub> | - | - |
| *P. aeruginosa* PA1 | Pus | NA | *bla*<sub>NDM-1</sub> | - | - |
| *P. aeruginosa* PA2 | Pus | NA | ND | - | - |
| *P. aeruginosa* PA3 | Pus | NA | ND | - | - |
| *P. aeruginosa* PA4 | Bronchial secretion | NA | ND | - | - |
| *P. aeruginosa* PA5 | Urine | NA | *bla*<sub>GES-1</sub> | - | - |
| *P. aeruginosa* PA6 | Unknown | NA | ND | - | - |
| *P. aeruginosa* PA7 | Blood | NA | *bla*<sub>GES-1</sub> | - | - |
| *P. aeruginosa* PA8 | Pus | NA | ND | - | - |
| *P. aeruginosa* PA10 | Pus | NA | ND | - | - |
| *S. typhi* ST1 | Blood | NA | ND | - | - |
| *S. typhi* ST2 | Unknown | NA | ND | - | - |
| *S. typhi* ST3 | Urine | NA | ND | - | - |
| *S. typhi* ST4 | Blood | NA | ND | - | - |
| *S. typhi* ST5 | Urine | NA | ND | - | - |
| *S. typhi* ST6 | Blood | NA | ND | - | - |
| *S. typhi* ST7 | Blood | NA | ND | - | - |
| *S. typhi* ST8 | Unknown | NA | ND | - | - |
| *E. cloacae* EL1 | Urine | NA | ND | - | - |
| *E. cloacae* EL2 | Blood | NA | *bla*<sub>NDM-1</sub> | - | - |
| *E. cloacae* EL3 | Urine | NA | *bla*<sub>NDM-1</sub> | *IncFIIA | - |
| *E. cloacae* EL4 | Bronchial secretion | NA | ND | - | - |
| *E. cloacae* EL5 | Blood | NA | *bla*<sub>OXA-181</sub> | - | - |
| *E. cloacae* EL6 | Urine | NA | - | - | - |
| *E. cloacae* EL7 | Urine | NA | *bla*<sub>IMP-1</sub> | - | - |
| *E. cloacae* EL8 | Urine | NA | *bla*<sub>GES-9</sub> | - | - |
| *A. baumannii* AB1 | Cerebrospinal fluid | NA | *bla*<sub>OXA-181</sub>,<br>*bla*<sub>OXA-51</sub> | - | - |
| *A. baumannii* AB2 | Urine | NA | *bla*<sub>OXA-181</sub>,<br>*bla*<sub>OXA-51</sub> | - | - |
| *A. baumannii* AB3 | Unknown | NA | *bla*<sub>OXA-51</sub> | - | - |
| *A. baumannii* AB4 | Pus | NA | *bla*<sub>OXA-23</sub>,<br>*bla*<sub>OXA-51</sub> | - | - |
| *A. baumannii* AB5 | Blood | NA | *bla*<sub>OXA-23</sub>,<br>*bla*<sub>OXA-51</sub> | - | - |
| *A. baumannii* AB6 | Urine | NA | *bla*<sub>OXA-51</sub> | - | - |
| *A. baumannii* AB7 | Urine | NA | *bla*<sub>OXA-51</sub> | - | - |
| *S. marcescens* SM1 | Bronchial secretion | NA | ND | - | - |
| *S. marcescens* SM2 | Blood | NA | *bla*<sub>NDM-1</sub> | - | - |
| *S. marcescens* SM3 | Unknown | NA | ND | - | - |
| *S. marcescens* SM4 | Urine | NA | ND | - | - |
| *S. marcescens* SM5 | Unknown | NA | ND | - | - |
| *A. xylosoxidans* AY1 | Unknown | NA | ND | - | - |
| *A. xylosoxidans* AY2 | Blood | NA | ND | - | - |
| *A. xylosoxidans* AY3 | Urine | NA | ND | - | - |
| *A. xylosoxidans* AY4 | Urine | NA | ND | - | - |
| *A. xylosoxidans* AY5 | Urine | NA | *bla*<sub>NDM-1</sub> | - | - |
| *K. oxytoca* KO1 | Blood | NA | ND | - | - |
| *K. oxytoca* KO2 | Urine | NA | ND | - | - |
| *K. oxytoca* KO3 | Blood | NA | ND | - | - |
| *K. oxytoca* KO4 | Blood | NA | *bla*<sub>OXA-181</sub> | - | - |
| *K. oxytoca* KO5 | Urine | NA | ND | - | - |
| *P. mirabilis* PM1 | Unknown | NA | ND | - | - |
| *P. mirabilis* PM2 | Blood | NA | *bla*<sub>OXA-181</sub> | - | - |
| *P. mirabilis* PM3 | Urine | NA | ND | - | - |
| *P. mirabilis* PM4 | Blood | NA | ND | - | - |
| *P. mirabilis* PM5 | Urine | NA | *bla*<sub>IMP-1</sub> | *IncFIA-FIB | - |
| *E. meningoseptica*<br>EM1 | Cerebrospinal<br>fluid | NA | ND | - | - |

NA- serotyping/pathotypic not applicable; ND- resistance gene not detected; \*Plasmids carrying resistance genes; '-' – Absence; '+' – Conjugation positive; highlighted in grey represents the isolates carrying resistance genes on conjugative plasmids

*K. pneumoniae* isolates harboured *bla*_NDM-1_ genes in IncFIA-FIB (KP10) and *bla*_GES-1_, *bla*_OXA-23/51-like_ genes in IncA/C (KP31 and KP39) type plasmids. One *E. cloacae* isolate harboured *bla*_NDM-1_ gene in IncFIIA (EL3) type plasmid and one *P. mirabilis* isolate harboured *bla*_IMP-1_ gene in IncFIA-FIB (PM5) type plasmid.

Overall, 6 *E. coli* (EC10, 21, 29, 39, 44, 47) isolates, 3 *K. pneumoniae* isolates (KP10, 31, 39), one *E. cloacae* isolate (EL3) and one *P. mirabilis* isolate (PM5) carried resistance genes on plasmid of the identified inc/rep types. All the 6 *E. coli* isolates (EC10, 21, 29, 39, 44, 47) were found to transfer resistance plasmids to susceptible *E. coli* AB1157. Inter-generic transfer of NDM-1 was observed in one *K. pneumoniae* isolate (KP10) in which *bla*_NDM-1_ harbouring plasmid IncFIA-FIB was transferrable to *E. coli* AB1157 (Table 4).

## 4.0 Discussion

In India, the prevalence of carbapenem-resistant Gram-negative bacteria has been reported with an increasing frequency [17,18]. In this study, the distribution of carbapenem-resistant isolates among 11 genera of Gram-negative bacteria isolated from diagnostic center in Tamil Nadu, India is reported. Previously, the increasing prevalence of ESBL and MBL producers among Gram-negative bacteria has been reported in India [27–30].

In this study, MIC results showed that 107/151 (71%) were resistant to meropenem in accordance with 128 isolates by the disk-diffusion method. All the 71 isolates harbouring carbapenem resistance genes were resistant with MIC and disc-diffusion method. The studied *E. coli* pathotypes (EPEC, EHEC, EIEC, EAEC and ETEC) are associated with intestinal diseases, they are collectively called as diarrheagenic *E. coli* (DEC) or intestinal pathogenic *E. coli* (IPEC) [31,32]. All these pathotypes are linked directly to their virulence properties and severity of infections. Though there are studies that showed the prevalence of DEC in India [33,34], still the studies on *E. coli* pathotypes (virulence) interrelation to carbapenem resistance is not well established in India. In this study, all the five *E. coli* pathotypes were found to harbour carbapenem resistance genes, namely EPEC (NDM-1, OXA-181, and OXA-23), EHEC (NDM-1, OXA-181, GES-1, and GES-9), EIEC (NDM-1, OXA-181), EAEC (GES-9, OXA-23, and GES-1), ETEC (NDM-1) and some are unknown pathotypes (Table 4). Adding to their virulence, the presence of resistance genes makes these bacterial infections (mostly diarrhoea) more complicated due to unavailability of treatment options. The *Klebsiella* isolates can be grouped into serotypes using surface antigens or surface exposed lipopolysaccharides [35]. The *Klebsiella* belonging to K-serotypes have K-antigen that relates to the capsule polysaccharide (CPS) [35]. Of the known capsular types (eight serotypes), the serotypes K1 and K2 are the most virulent among the hypervirulent *K. pneumoniae* (hvKP) [36]. In this study, of the 14 *Klebsiella* isolates belonging to K1 serotypes, six isolates carried carbapenem resistance genes, *bla*_NDM-1_, *bla*_OXA-181_ and *bla*_OXA-51_. Among the 11 *Klebsiella* isolates belonging to K2 serotype, two isolates were found to carry carbapenem resistance genes, *bla*_OXA-181_ and *bla*_GES-9_ (Table 4). In India, NDM-1 and OXA-48 genes were detected in *Klebsiella* belonging to K2 serotypes [36], but to the best of our knowledge, this is the first study to detect the presence of carbapenemase genes NDM-1, OXA-181 and OXA-51 among K1 serotypes.

As carbapenems are the last resort of antibiotics available to treat infections caused by Gram-negative bacteria, the prevalence of carbapenem resistance is given a global attention. Our previous studies had reported the dissemination of carbapenem-resistant bacteria and carbapenem resistance genes among Gram-negative bacteria [17,18]. Here, we report the prevalence (71%) of carbapenem resistant isolates among 11 genera of Gram-negative bacteria. Beta-lactamase resistance genes such as *bla*_NDM-1_ (n=22), *bla*_OXA-181_ (n=21), *bla*_GES-1_ (n=11), *bla*_GES-9_ (n=8), *bla*_OXA-23_ (n=7), *bla*_OXA-51_ (n=9) and *bla*_IMP-1_ (n=3) were found in 71 isolates (10 isolates carrying more than one genes), comparatively our earlier studies showed the low prevalence (27%) of *bla*_NDM-1_ and *bla*_OXA-181_ genes among carbapenem-resistant isolates [18]. The coexistence of *bla*_NDM-1_ and *bla*_OXA-181_ in *E. coli* is one of the serious concerns from healthcare prospective. All the *A. baumannii* isolates (n=7) were found to have either the class D carbapenem hydrolyzing oxacillinases (OXA-23, OXA-181) and OXA-51 is naturally existed in *Acinetobacter* spp. [37]. There were earlier reports in India showing the presence of OXA-23 and OXA-51 in carbapenem-resistant *Acinetobacter* causing serious health care problems [12]. *Enterobacteriaceae* are encoded by OXA-48-like genes as carbapenem-hydrolyzing class D β-lactamases [13,38]. But the unusual occurrence of *bla*_OXA-23_ in *E. coli*, and plasmid-borne (IncA/C) *bla*_OXA-23_ and *bla*_OXA-51_ in *K. pneumoniae* is one of the important findings of this study. There are very few earlier studies that reported the presence of *bla*_OXA-23_ gene in *E. coli* [39,40]. To the best of our knowledge, this is the first study to report the plasmid-borne (IncA/C) OXA-23 and OXA-51 in *K. pneumoniae*. The OXA-23-like genes in *Enterobacteriaceae* may be carried within a transposon but was not characterized in this study. The resistance reports on *E. meningoseptica* are very rare in India [39,40] and in our study, it was found that one isolate of *E. meningoseptica* was resistant to imipenem and meropenem. Though earlier studies showed the presence of carbapenemase genes in *E. meningoseptica*, in this study no carbapenem resistance genes were amplified.

Carbapenem resistance among Gram-negative bacteria is becoming very common in India and the spread of carbapenem resistance genes are one of the troublesome problems. These resistance genes that are located adjacent to the mobile genetic elements (integrons and transposons), which facilitates the easy transposition between replicons [41]. The most common plasmid replicon types for carbapenem resistance genes are IncF, IncA/C_2_, IncX3, IncL/M and IncH [42]. In this study, *bla*_NDM-1_ was found to be harboured in IncX, IncA/C, IncFIA-FIB and IncFIIA; *bla*_OXA-181_ in IncA/C, IncFIA-FIB and IncFIIA; *bla*_GES-1/9_ in IncFIA-FIB and IncA/C; *bla*_IMP-1_ in IncFIA-FIB and *bla*_OXA-23/51_ in IncA/C. The presence of plasmid-borne *bla*_OXA-23/51_ is very rare and important finding, considering the rapid spread of carbapenem-resistance among Gram-negative bacteria. Interestingly, the isolates such as *P. aeruginosa*, *Salmonella typhi*, *A. baumannii*, *S. marcescens*, *A. xylosoxidans*, *K. oxytoca*, and *E. meningoseptica* do not carry any plasmids harbouring resistance genes. This clearly showed that the beta-lactamase or carbapenemase resistance genes were present in the plasmids with different replicon types in the study region. Earlier, the *bla*_NDM_ IncFII plasmids were reported from India [42] and IncFIA-FIB plasmids carrying carbapenem resistance genes such as *bla*_NDM_ was report from India in the samples collected from river and sewage treatment plants [42,43]. This study also showed that some plasmids were carrying more than one resistance genes which are an alarming threat to the public health. Conjugative plasmids are known to spread their resistance characteristics among the bacteria from the same or different genus. This study showed that all the six *E. coli* isolates carrying plasmid-borne resistance genes (*bla*_NDM-1_, *bla*_OXA-181_, *bla*_GES-1_, *bla*_GES-9_) were conjugative and one *K. pneumoniae* isolate plasmid (IncFIA-FIB with *bla*_NDM-1_) was transferable which clearly shows the way by which resistance genes can rapidly spread in clinical bacteria.

## 5.0 Conclusion

The emerging antibiotic resistance in bacteria is a worrisome problem. This study highlighted the distribution of carbapenem resistant isolates in the study region with the extra emphasis on the existence of *bla*_NDM-1_, *bla*_OXA-48-like_, *bla*_IMP-1_, *bla*_GES-1_, *bla*_GES-9_, *bla*_OXA-23-like_, *bla*_OXA-51-like_ among the clinical pathogens. Alternative therapeutic options should be undertaken immediately to combat the problem of resistance especially to treat infections caused by carbapenem resistant bacteria. Our study shows that the ‘conjugative plasmids’ can strongly contribute to the resistance transfer in pathogens leading to dissemination of resistance genes. Alternative approaches are necessary to combat the problem of resistance and concepts such as ‘one-health approach’ can be appreciated.

## Authors’ contribution

Authors PM and NR, collected the isolates from the clinical samples. Authors PM and NR undertook the laboratory work, NR and BSL interpreted the data, and PM and NR wrote the initial manuscript. Authors NR and BSL revised and edited the manuscript. All the authors’ have read and approved the manuscript.

## Ethics approval

Ethical approval from Institutional Ethical Committee for studies on Human subjects (IECH), ref. no. VIT/IECH/004/Jan2015

## Availability of data and materials

All the datasets are presented in the main manuscript. The raw datasets used and/or analysed during the current study are available from the corresponding author on reasonable request.

## Acknowledgement

The authors would like to thank Vellore Institute of Technology (VIT) for providing partial funding, ‘VIT Seed Grant’ and Council of Scientific and Industrial Research (CSIR) for providing financial assistance to PM in the form of senior research fellowship (SRF) to support this research.

